# Identification of a polymorphism in the N gene of SARS-CoV-2 that adversely impacts detection by a widely-used RT-PCR assay

**DOI:** 10.1101/2020.08.25.265074

**Authors:** Manu Vanaerschot, Sabrina A. Mann, James T. Webber, Jack Kamm, Sidney M. Bell, John Bell, Si Noon Hong, Minh Phuong Nguyen, Lienna Y. Chan, Karan D. Bhatt, Michelle Tan, Angela M. Detweiler, Alex Espinosa, Wesley Wu, Joshua Batson, David Dynerman, CLIAHUB Consortium, Debra A. Wadford, Andreas S. Puschnik, Norma Neff, Vida Ahyong, Steve Miller, Patrick Ayscue, Cristina M. Tato, Simon Paul, Amy Kistler, Joseph L. DeRisi, Emily D. Crawford

## Abstract

We identify a mutation in the N gene of SARS-CoV-2 that adversely affects annealing of a commonly used RT-PCR primer; epidemiologic evidence suggests the virus retains pathogenicity and competence for spread. This reinforces the importance of using multiple targets, preferably in at least 2 genes, for robust SARS-CoV-2 detection.

**Article Summary Line:** A SARS-CoV-2 variant that occurs worldwide and has spread in California significantly affects diagnostic sensitivity of an N gene assay, highlighting the need to employ multiple viral targets for detection.

## Introduction

Standard diagnostic testing for SARS-CoV-2 infection involves examination of respiratory specimens for the viral genome by RT-PCR. Typically, this is done with multiple primer pairs targeting more than one viral gene. However, recent WHO guidelines (*1*) state that “In areas where the COVID-19 virus is widely spread a simpler algorithm might be adopted in which, for example, screening by RT-PCR of a single discriminatory target is considered sufficient.” Currently, 36 out of 175 FDA-approved Emergency Use Authorizations involve assays that employ just a single target for RT-PCR (*2*).

Here we report the identification of a nucleotide change in an N gene primer sequence that impairs annealing and amplification, resulting in increased C_t_ values and decreased diagnostic sensitivity. Our results indicate the potential hazard of relying on one target when assaying for SARS-CoV-2, even in areas of high prevalence.

### The Study

Since April 7, 2020, the COVID-19 diagnostic laboratory at the Chan Zuckerberg Biohub and the University of California San Francisco (CLIAHUB) has been receiving samples for no-cost testing from multiple counties in California. Our real-time RT-PCR protocol (*3*) employs N gene (NIID_2019-nCov_N_F2/R2ver3/P2 (*4*)) and E gene (E_Sarbeco_F/R/P1 (*5*)) simplex assays that have been used widely by other groups. In July 2020, we identified geospatial and temporal clustering of 35 samples from Madera County, California that demonstrated poor N gene assay performance relative to the E gene assay. We initiated an investigation to identify possible causes.

Figure 1A shows the concordance of C_t_ values for the two assays in the 3,957 positive tests conducted in CLIAHUB during May 27 — August 7, 2020. For samples with positive E and N gene results (n=3629), the N and E gene C_t_ value difference (ΔC_t_(N-E)) was 0.40 ± 1.18 (mean ± standard deviation). We defined potential mutant sample set A as those with an ΔC_t_(N-E) ≥2.96 (2.5 standard deviations above the mean) and with an E C_t_ value ≤30. The ΔC_t_(N-E) in sample set A was on average 5.35, implying an ~41-fold impairment of N gene amplification. All 42 samples were collected during June 30 — August 4, and 34 originated from Madera County. We defined sample set B as samples from Madera County processed during the same time period with an E C_t_ >30 and ΔC_t_(N-E) >2.96 or an E C_t_ >30 and for which no N C_t_ was detected. Importantly, the N C_t_ values of these 19 samples in sample set B, if detected, often exceeded the C_t_ corresponding to the limit of detection of 100 viral genome copies per ml, affecting the accuracy of these C_t_ values.

**Figure 1.**
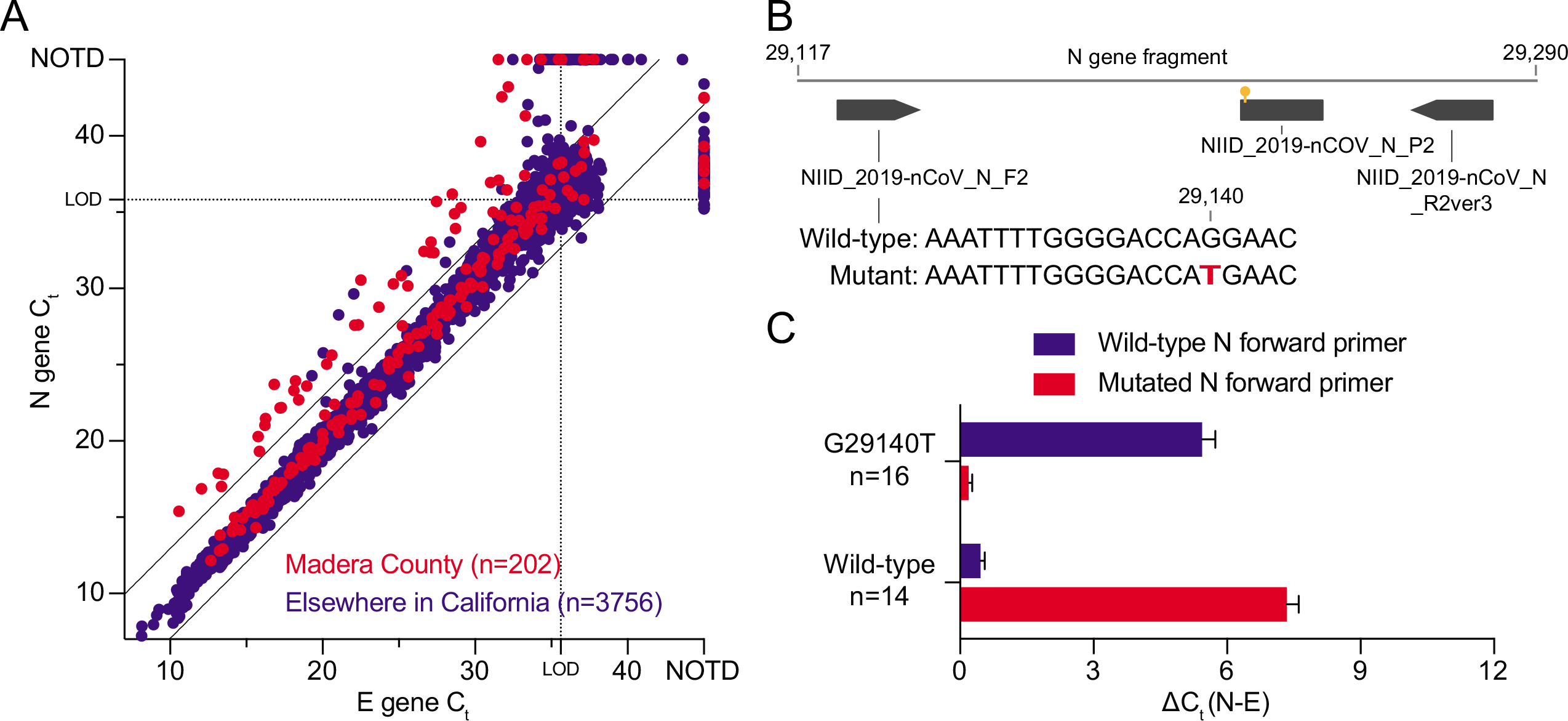
Single mutation in forward N gene primer binding site decreased SARS-CoV-2 RT-PCR sensitivity. (A) Potential SARS-CoV-2 mutants were identified by their increased ΔC_t_ between the N and E gene assays (>2.5 □ standard deviation of average ΔC_t_, cut-off indicated by black lines). Dotted lines indicate the average C_t_ value at the limit of detection (LOD) of each assay, above which more variation is expected. NOTD: not detected. Raw data is available in Appendix Table 1. (B) Diagram showing a fragment of the N gene, with the N gene primers and probe originally developed by the National Institute of Infectious Diseases (NIID) in Tokyo, Japan (*4*) and the identified G29140T mutation indicated. (C) The increased ΔC_t_(N-E) of mutant lines using the conventional RT-PCR with wild-type primer was reversed when a primer incorporating the mutation was used. The opposite was observed for wild-type samples that showed an increased ΔC_t_(N-E) when the mutated primer was used, further validating causality of the G29140T mutation for reduced N gene RT-PCR performance. Error bars indicate the standard error of the mean. Raw data is available in Appendix Table 2.

Sequencing of the detected N gene fragment for the 42 potential mutant samples from set A revealed a single mutation in the forward primer binding site (16^th^ of 20 nucleotides) in the N gene, corresponding to G29140T in the SARS-CoV-2 genome, in 34 samples (33 from Madera County, 1 from the Bay Area). The remaining 8 samples showed a wild-type N gene fragment, suggesting that the higher average ΔC_t_(N-E) of 5.99 in these samples was an artefact. None of the 17 randomly-selected control samples from Madera with a ΔC_t_(N-E)<2.96 showed G29140T. Interestingly, a G29135A mutation (11^th^ of 20 nucleotides of the forward primer) was observed in a single sample that showed a ΔC_t_(N-E) of 2.33, indicating a more modest effect compared to the G29140T mutation.

For potential mutant sample set B, sequences were obtained for 14 out of 19 samples. Twelve of these carried the G29140T mutation. Importantly, the N gene RT-PCR did not detect SARS-COV-2 in 5 of these 10 samples. Fortunately, these cases could still be recognized as infected by use of the E gene assay.

To further investigate the effect of the G29140T mutation on N gene PCR performance, we synthesized a new N gene forward primer with full complementarity to the mutant sequence and compared this mutated primer to the canonical primer for a subset of 16 mutant and 14 wild-type samples (Figure 1C). For mutant samples, the ΔC_t_(N-E) dropped from 5.44 with the canonical primer to 0.19 with the mutated primer. This trend was inverted for wild-type samples where the ΔC_t_(N-E) increased from 0.46 with the canonical primer to 7.34 with the mutated primer. These data validate that the G29140T mutation is causative for the observed aberrant N gene C_t_ values.

The G29140T mutation encodes a Q289H amino acid mutation in the N gene that was also found in other sequences available on GISAID (*6*). Sequences with an unambiguous Q289H mutation originated from England (n=14), the United States (n=5, New York, Utah, California, Michigan and Texas), the Democratic Republic of Congo (n=2), Iceland (n=2), Sweden (n=2), Japan (n=1), and Scotland (n=1). Three others from Austria, Australia, and Switzerland showed ambiguous calls possibly resulting in Q289H. Other mutations at the same position (Q289K, Q289L, Q289R and Q289*) are singular occurrences in GISAID for which artefacts could not be ruled out. Amino acid 289 is located within the dimerization domain of the nucleocapsid protein but is not involved in any known dimer interface interactions, though tertiary structure level interactions could be impacted by mutations at this position (*7*).

Whole genome sequencing of 20 mutant samples and 11 wild-type samples from Madera County showed that all mutant samples clustered together, displaying little genetic diversity (Figure 2). Of the 27 GISAID sequences sharing the unambiguous Q289H variant (*6*), one from San Diego County was identical by descent to the Madera cluster. The remaining 26 fell on different clades of the tree, with 11 estimated recurrent mutation events at the locus. Furthermore, 6 of these were identical at the amino acid level, but carried a different mutation at the nucleotide level (G29140C instead of G29140T). Among the wild-type samples from Madera, one was a close relative to the mutant cluster, sharing a common ancestor shortly before the mutation arose.

Madera County is currently experiencing significant community spread of COVID-19, with a 14-day incidence rate of 378 cases per 100,000 population during July 1 — July 14, when the first mutant sample was observed (*8*). Out of 202 total samples from this county, 45 (22.3%) carried the G29140T mutation. These originated from four of thirteen mobile community testing sites in Madera County that covered both residential and also rural areas where primarily agricultural workers reside, suggesting broad geographic distribution of cases within the county. Nine of the samples originated from a single outbreak at a skilled nursing facility. Limited case investigation data from Madera County patients indicate that 17 of 19 individuals reporting symptom status did exhibit COVID-19 symptoms (89%) with two deaths (4.8%). Cases occurred in individuals in a mix of professions [out of 14 reporting: 1 healthcare worker (7%); 1 laborer (7%); 2 craftsman (14%); 2 retired (14%); 3 unemployed (21%); 4 farmworkers (29%)] and predominantly among Hispanic or Latinx persons [13/18 (72%)].

**Figure 2.**
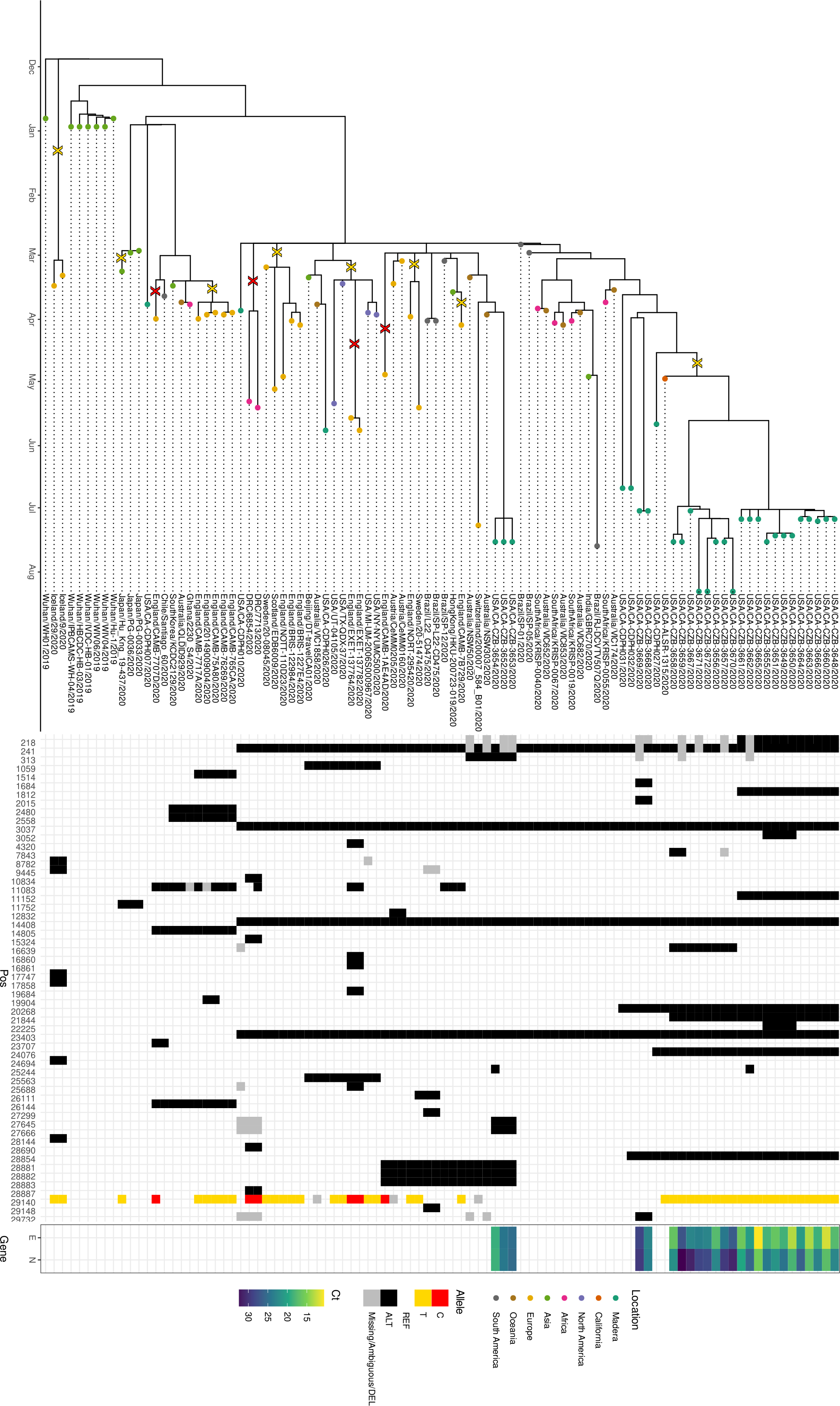
Phylogeny of SARS-CoV-2 isolates with N mutation. Tree, haplotype map, and C_t_ values for SARS-CoV-2 isolates, including those with the G29140T mutation. Inferred mutation events on the tree are annotated with an X that is colored depending on the allele. Both synonymous variants of the Q289H mutant are found, with the mutation estimated to have recurred 11 times on the tree, with only one of the mutant samples from GISAID being identical by descent to the Madera cluster. One of the wild-type Madera samples was closely related to the mutant cluster, with a common ancestor just before the mutation event. SARS-CoV-2 genome sequences from Madera samples were recovered via Primal-Seq Nextera XT version 2.0 (modified from (*9*)), with the ARTIC Network v3 primers (*10*)), followed by paired-end 150bp sequencing on the Illumina platform. Reads were aligned to the reference genome (genbank accession MN908947.3) with minimap2 (*11*), samtools (*12*) to generate a pileup, and ivar (*13*) to trim primers. A phylogenetic tree was estimated using Nextstrain’s ncov repository (*14*). Additional samples from GISAID were selected by searching CoV-GLUE for sequences with the Q289H mutation, and by using Nextstrain’s sparse subsampling scheme to select additional wildtype samples for context. Sequence data downloaded from GISAID is available in Appendix Table 3, all code used for analyses and figure generation is publicly available (*15*).

## Conclusions

Our data show that even in areas of high SARS-CoV-2 community spread, replication-competent mutations can arise that impair RT-PCR primer recognition and could lead to reduced test sensitivity and under-diagnosis if labs adopted the practice of employing only one target for SARS-CoV-2 detection. Our findings strongly support the routine use of at least two targets for SARS-CoV-2 detection by RT-PCR.

## Acknowledgements

We thank Peter Kim and Don Ganem for helpful discussions and editorial assistance on the manuscript. This study was supported by the Chan Zuckerberg Biohub.

## Data and code availability

All code and C_t_ data are available in our Github repository: https://github.com/czbiohub/polymorphism_sarscov2_diagnostics. RT-PCR for all samples is available in Appendix Table 1. RT-PCR data with the mutated primer is available in Appendix Table 2. Sequence data is available via GISAID; see Appendix Table 3.

## Author Bio

Dr. Manu Vanaerschot is a scientist at the Chan Zuckerberg Biohub, San Francisco, California, USA. His primary research interests are early detection and management of infectious disease outbreaks.

Appendix Table 1. Raw data detailing N and E gene C_t_ values and location data for samples depicted in Figure 1A.

Appendix Table 2. Raw data detailing the individual ΔC_t_(N-E) data points depicted in Figure 1C.

Appendix Table 3. Metadata of sequences depicted in Figure 2.

